# Gene isoforms as expression-based biomarkers predictive of drug response *in vitro*

**DOI:** 10.1101/160937

**Authors:** Zhaleh Safikhani, Kelsie L. Thu, Jennifer Silvester, Petr Smirnov, Mathieu Lupien, Tak W. Mak, David Cescon, Benjamin Haibe-Kains

## Abstract

**Background:** One of the main challenges in precision medicine is the identification of molecular features associated to drug response to provide clinicians with tools to select the best therapy for each individual cancer patient. The recent adoption of next-generation sequencing technologies enables accurate profiling of not only gene expression but also alternatively-spliced transcripts in large-scale pharmacogenomic studies. Given that altered mRNA splicing has been shown to be prominent in cancers, linking this feature to drug response will open new avenues of research in biomarker discovery.

**Methods:** To address the lack of reproducibility of drug sensitivity measurements across studies, we developed a meta-analytical framework combining the pharmacological data generated within the Cancer Cell Line Encyclopedia (CCLE) and the Genomics of Drug Sensitivity in Cancer (GDSC). Predictive models are fitted with CCLE RNA-seq data as predictor variables, controlled for tissue type, and combined GDSC and CCLE drug sensitivity values as dependent variables.

**Results:** We first validated the biomarkers identified from GDSC and CCLE using an existing pharmacogenomic dataset of 70 breast cancer cell lines. We further selected four drugs with the most promising biomarkers to test whether their predictive value is robust to change in pharmacological assay. We successfully validated 10 isoform-based biomarkers predictive of drug response in breast cancer, including TGFA-001 for the MEK tyrosine kinase inhibitor (TKI) AZD6244, DUOX-001 for the EGFR inhibitor erlotinib, and CPEB4-001 transcript expression associated with lack of sensitivity to paclitaxel.

**Conclusion:** The results of our meta-analysis of pharmacogenomic data suggest that isoforms represent a rich resource for biomarkers predictive of response to chemo- and targeted therapies. Our study also showed that the validation rate for this type of biomarkers is low (<50%) for most drugs, supporting the requirements for independent datasets to identify reproducible predictors of response to anticancer drugs.

## INTRODUCTION

Cell lines are the most widely-used cancer models to study response of tumors to anticancer drugs. Not only have these cell lines recently been comprehensively profiled at the molecular level, but they have also been used in high-throughput drug screening studies, such as the Genomics of Drug Sensitivity in Cancer (GDSC) [1] and the Cancer Cell Line Encyclopedia [2]. The overarching goal of these seminal studies was to identify molecular features predictive of drug response (predictive biomarkers). Consequently, the GDSC and CCLE investigators were able to confirm a number of established gene-drug associations, including association of ERBB2 amplification with sensitivity to lapatinib and BCR/ABL fusion expression and nilotinib. They also found new associations such as SLFN11 expression and response to topoisomerase inhibitors, thereby supporting the potential relevance of cell-based high-throughput drug screening for biomarker discovery. However the biomarkers validated in preclinical settings are still largely dominated by genetic (mutation, copy number alteration or translocation) as opposed to transcriptomic (gene expression) features. Therefore, there is a need for further investigation of transcriptomic markers associated with drug response in cancer.

The vast majority of pharmacogenomic studies investigated the association between gene-specific mRNA abundance and drug sensitivity [1–6]. However, it is well established that genes undergo alternative splicing in human tissues (61% of the genome; Ensembl version 37), and changes in splicing have been associated with all hallmarks of cancer [7]. Despite the major role of alternative splicing in cancer progression and metastasis [7], only a few small-scale studies have reported associations between these spliced transcripts (also referred to as isoforms) and drug response or resistance [8–10]. These limited, yet promising associations support the potential relevance of isoform expression as a new class of biomarkers predictive of drug response. Among the mRNA expression profiling technologies, high-throughput RNA sequencing (RNA-seq) enables quantification of both isoform and gene expression abundances at the genome-wide level. Recent studies have highlighted the advantages of RNA-seq over microarray-based gene expression assays [11–15]. In particular, microarray profiling platforms are limited to pre-designed cDNA probes [11] and they depend on background levels of hybridization. They also suffer from limited dynamic range probe hybridization. Since the detection of transcripts and genes using RNA-seq is based on high resolution short reads sequencing instead of probe design, they have the potential to overcome these limitations [13].

Recent initiatives have profiled hundreds of cancer cell lines using Illumina RNA-seq technology [3,16–18]. As part of CCLE, the Broad Institute of Harvard and MIT recently released RNA--seq profiles of 935 cancer cell lines through the Cancer Genomics Hub (CGHub) [19]. Two other initiatives used RNA-seq to profile panels of 70 (GRAY [3]) and 84 (UHN [17]) breast cancer cell lines. The availability of these valuable datasets offers unprecedented opportunities to further explore the transcriptomic features of cancer cells and study their association with drug response. Here, we explore the genome-wide transcriptomic landscape of large panels of cancer cell lines to identify isoform-level expression features predictive of drug response *in vitro*. Based on our new meta--analytical framework combining the GDSC and CCLE drug sensitivity data for biomarker discovery, we show that isoform-level expression measurements are more predictive of response to cytotoxic and targeted therapies than are gene-level expression values. We tested the accuracy of our most promising isoform biomarkers in two independent breast cancer pharmacogenomic datasets, GRAY and UHN. We validated ten isoform-based biomarkers predictive of response to lapatinib, erlotinib, AZD6244 (MEK inhibitor) and paclitaxel, indicating that isoforms constitute a promising new class of biomarkers for cytotoxic and targeted anticancer therapies.

## MATERIALS AND METHODS

A schematic view of the design of our study is shown in Figure 1.

**Figure 1:**
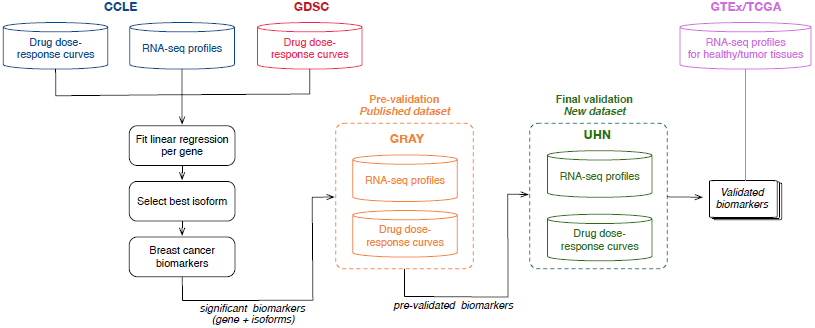
Analysis design of the study. CCLE (in blue) and GDSC (in red) are used to identify a set of biomarkers significantly associated with response to each of the 15 drugs screened in both training sets. The biomarkers predictive in breast cancer cell lines are selected and further validated in an independent, *in vitro* breast cancer dataset (GRAY). This step, referred to as pre-validation, enables the selection of generalizable, isoform-based biomarkers for breast cancer (represented in orange). The newly generated UHN dataset is then used to test whether the selected isoform-based biomarkers are robust to the use of a different pharmacological assay (final validation represented in green). The expression distribution of the final set of biomarkers is compared between patient tumors (TCGA) and healthy tissues (GTEx).

### Published Pharmacogenomics studies

We used our *PharmacoGx* platform [20] to create curated, annotated and standardized pharmacogenomic datasets composed of CCLE [2], GDSC [1] and GRAY [3]. CCLE and GRAY pharmacological data were generated using the CellTiter-Glo assay (which quantitates ATP, Promega), while GDSC used the Syto60 assay (a nucleic acid stain, Invitrogen) [21]. We updated CCLE and GRAY PharmacoSets to include gene and isoform-level expression data processed from the raw RNA-seq profiles downloaded from CGHub [19] and NCBI GEO [22], respectively.

### RNA-seq data processing

We used Tophat2 [24] using the EnsemblGenome Reference Consortium release GRCh37 [25]. Cufflinks [26] is used to annotate genes and isoforms and quantify their expression. Gencode version 12 [27] was used as the transcript model reference for the alignment as well as for all gene and isoform quantifications. Gencode annotated a total of 53,934 genes, which includes 20,110 protein coding genes, 11,790 long noncoding RNA’s (lncRNA’s), and 12,648 pseudogenes. Expression values were computed as the log_2_(FPKM+1) where FPKM represents the number of fragments per kilobase per million mapped reads units which control for sequence length and sequencing depth [28].

### Pharmacological data processing

We developed a unified framework to process the raw pharmacological data of CCLE, GDSC and GRAY and to obtain the drug dose-response curves using a standard curve fitting algorithm [20] (Supplementary Methods). To summarize the drug dose-response curves into a single sensitivity measure we computed the area under the curve (AUC) metric, which combines both potency and efficacy of drug responses [29] (Supplementary Figure 1; Supplementary Methods). Compared with IC_50_ and E_max_ metrics, which represent only one point on the drug dose-response curve, AUC values are computed by integrating all data points. Consequently, AUC has been shown to be more reproducible across pharmacogenomic studies [30,31]. In this study, we used the area above the drug dose-response curve (AAC=1-AUC; Supplementary Figure 1) so that higher AAC represent high drug sensitivity.

### Biomarker discovery

To identify gene and isoform expression robustly associated with drug sensitivity, we developed a machine learning pipeline combining linear regression models with a bootstrapping procedure for stringent model selection. Our choice of model assumes a linear relationship between molecular features and drug responses. Although violation of this assumption may result in biased predictions, linear models are robust to variation or noise in the data, making them less prone to overfitting in a high-dimensional context such as pharmacogenomics. Therefore the association between each molecular feature and response to a given drug is assessed by fitting linear models using the gene or isoform expression across cell lines as predictor variables, adjusted for tissue of origin of cancer cell lines, and their sensitivity values to the given drug as dependent variables (Supplementary Figure 2). To assess the association of each gene and its isoforms to a given drug, three linear models were constructed for each dataset as following.

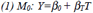

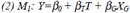

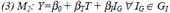

**Figure 2:**
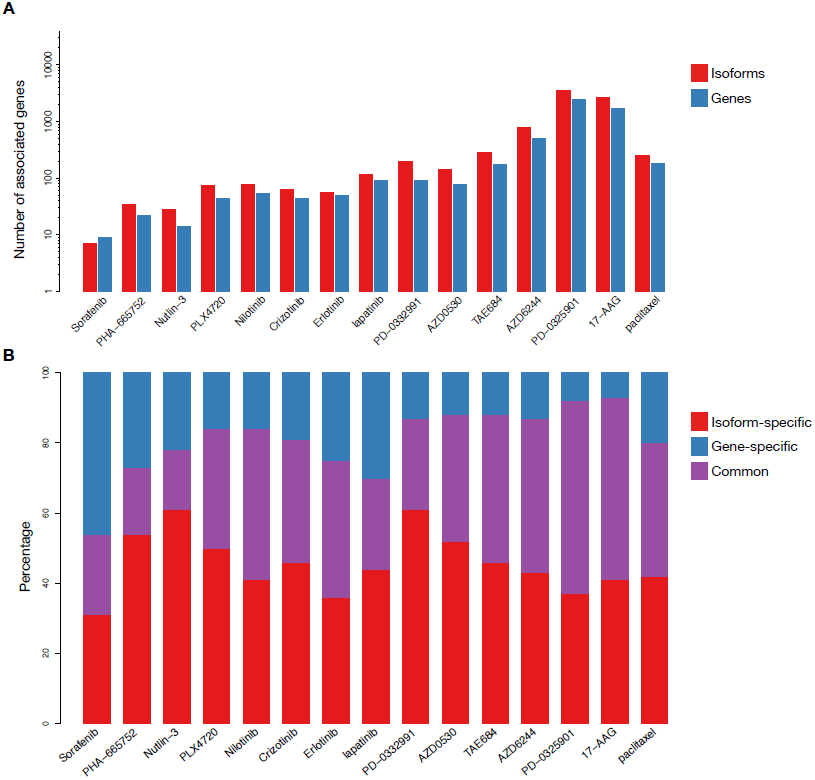
Comparison of number of statistically significant predictive biomarkers for each of the 15 drugs in common between CCLE and GDSC. (A) Number of significant biomarkers at the levels of gene and isoform expression. (B) Proportion of biomarkers that are significant at the gene level, isoform levels or both.

Where *T* represents the tissues of origin as a vector of size *N* × *1*; *N* is the number of cell lines; *Y* denotes the drug sensitivity vector of size *N* ×*1* containing the drug sensitivity values (AAC) of the cell lines treated by the drug of interest; *X*_*G*_ represents a vector of size *N* × *1* of log_2_ normalized FPKM values for the expression of gene *G* across all the cell lines; *G*_*I*_ is all the isoforms of gene *G*; *I*_*G*_ is a vector of size *N* × *1* of log_2_ normalized FPKM values for each isoform of *G* across all the cell lines. The effect size of each association is quantified by *β*_*G*_ and *β*_*I*_, which indicate the strength of associations between drug response and the molecular feature of interest, adjusted for tissue type. To estimate standardized coefficients from the linear model, the variables *X*_*G*_ and *I*_*G*_ are scaled (standard deviation equals to one, mean equals to zero). The null model (Equation (2)) estimates the association between drug response and tissue of origins. The models in Equations (3) and (4) estimate the strength and significance of the association between drug sensitivity and the gene-level and its best isoform expressions, respectively.

To address the lack of reproducibility of drug sensitivity measurements across studies [30,32], we developed a meta-analytical pipeline to combine the pharmacological data from CCLE and GDSC. The June 2014 release of CCLE consists of 11,670 experiments in which 24 drugs have been screened on 1,053 cancer cell lines from 24 tissue origins. GDSC release 5 comprises of 79,903 experiments for 140 different drugs tested on a panel of up to 778 unique cell lines from 30 tissue types. The panel of drugs and cell lines screened in these two datasets overlapped for 15 compounds and 512 cell lines, respectively (Supplementary Files 1 and 2, Supplementary Figure 3). Univariate gene-drug associations were computed using the linear models described in above-mentioned equations with CCLE RNA-seq data as predictors and CCLE and GDSC drug sensitivity data separately. We recognize that using CCLE RNA-seq data in combination with GDSC is suboptimal as gene expression of cell lines are subject to biological and technical variations [33]. In the absence of RNA-seq data for GDSC, we could only address the variations observed in the drug sensitivity measurements, which we demonstrated to be significantly higher than variations in gene expression data [32]. To ensure that cell line identity was conserved across CCLE and GDSC, we performed SNP fingerprinting (Supplementary Methods) and filtered out the cell lines identified as different across studies using a cutoff of 80% concordance [32]. In addition we compared the microarray expression profiles of cell lines between microarray and RNA-seq profiles, which resulted in good concordance (Supplementary Figure 4) supporting that expression profiling are consistent.

**Figure 3:**
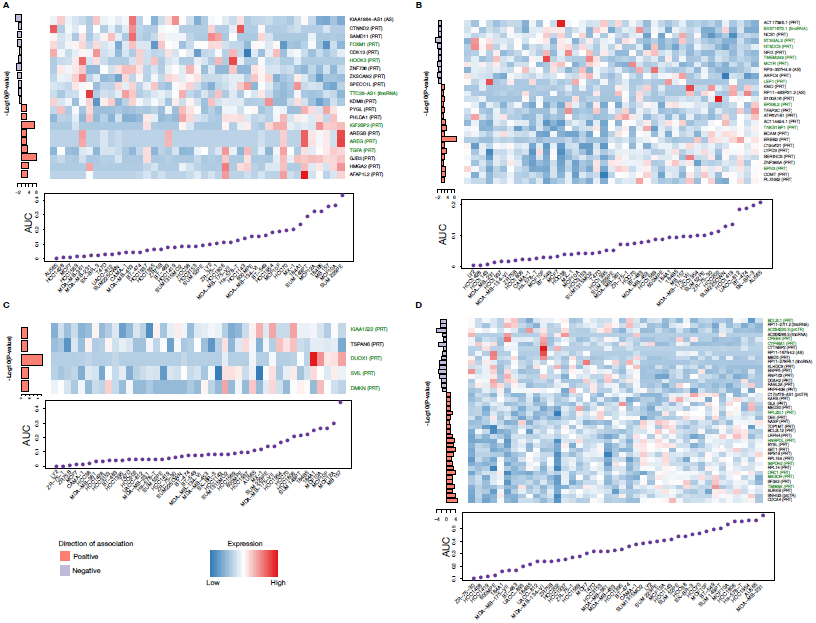
Isoform-based biomarkers successfully pre-validated in the independent GRAY dataset for (A) AZD6244, (B) lapatinib (C) erlotinib, and (D) paclitaxel. Cell lines are ordered by their sensitivity to the drug of interest and their isoform expression is shown in the heatmap, with the drug sensitivity (AUC) plotted below. The left side bar plot shows the significance of the association between isoform expression and drug sensitivity as the -log10(p-value) multiplied by the sign of the coefficient in the corresponding regression model. Genes for which the candidate isoform is significantly more predictive than its corresponding overall gene expression values are represented in green.

**Figure 4:**
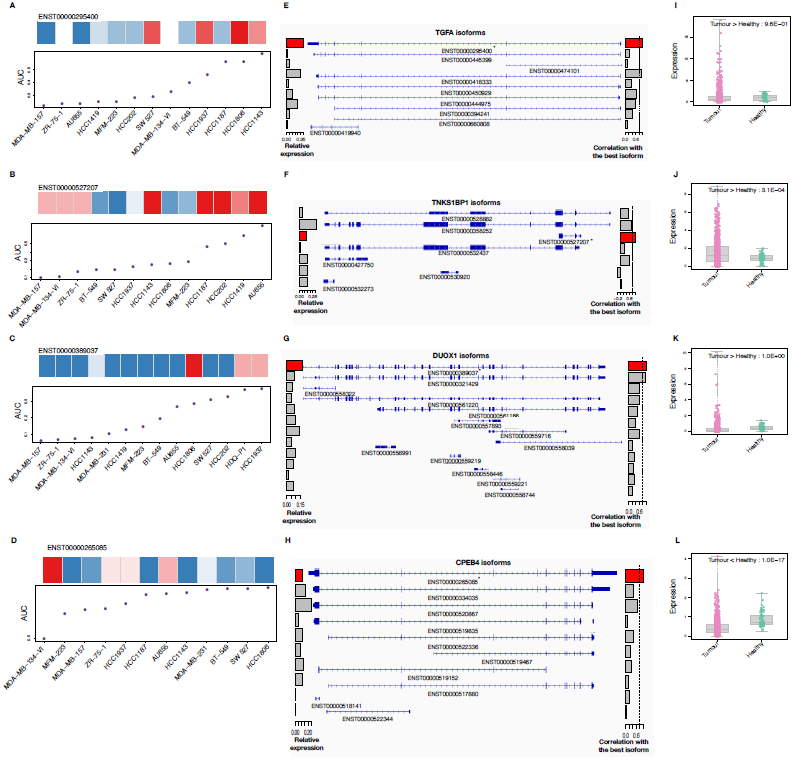
Validation of the candidate isoforms predictive of response to (A,E,I) AZD6244, (B,F,J) lapatinib (C,G,K) erlotinib, and (D,H,L) paclitaxel in the independent UHN dataset generated where a different pharmacological assay (sulforhodamine B assay) was used to measure drug sensitivity. In panels A-D, cell lines are ordered by their sensitivity to the drug of interest and their isoform expression is shown in the heatmap, with the drug sensitivity (AUC) plotted below. In panels E-H, exon occupancy of each candidate isoform (*) is visualized using the USCS Genome Browser, with a barplot on the right side representing the correlation (ρ) of expression between each isoform and the candidate isoform (red bar). A vertical dashed line represents ρ = 0.8 to identify highly correlated isoforms of the same gene. Panels I-L enables statistical comparison of the candidate isoform expression distribution across breast patient tumors and heathly tissues.

The predictive value (R^2^) and significance (p-value) of the fitted models are estimated using the linear models described in Equations (2) and (3). To determine the most predictive isoform for each gene the predictive value of all of its isoforms is estimated using equation (3) and the most significant isoform (the one with the smallest bonferroni-corrected p-value) is selected for further analysis. Comparison of the predictive value of each model was performed using a bootstrapping procedure: 100 resampled datasets are generated where the cell lines are obtained by sampling with replacements from all the cell lines with sensitivity and expression profile available for a given drug. The linear regressions are solved for each bootstrap using the resampled set (∼ 2/3) and unselected cell line set (∼ 1/3) for training and testing, respectively. To evaluate the prediction performance of a gene or isoform model, its vector of R^2^ values is compared to a null model using a one-sided wilcoxon signed rank test. Bootstrapping procedure is applied on the gene and its most predictive isoform. To combine the fitted models obtained from CCLE and GDSC, their coefficients and p-values were averaged and weighted by the number of cell lines in those datasets (Supplementary Figure 2). To control for multiple testing, we corrected the p-values obtained for all genes and isoforms, separately, using the false discovery rate (FDR) method [34].

### Pre-validation of isoform-based biomarkers (GRAY)

We validated the accuracy of our biomarkers using a previously-published independent dataset, GRAY [3], which includes RNA-seq of a panel of 70 breast cancer cell lines screened with 90 FDA-approved drugs (CellTiter-Glo pharmacological assay; Supplementary Table 1), with 8 compounds in common with CCLE and GDSC (Supplementary Figure 5). To check the predictive value of our biomarkers in breast cancer, we fitted the linear models in Equations (1) to (3) using only breast cancer cell lines in our training sets. A biomarker is selected if its predictive value in breast cancer cell lines is greater than or equal to the predictive value across all tissue types. To validate the selected biomarkers in GRAY we computed the significance of the linear association between the biomarker expression and drug response (p-value < 0.05) with the same direction of association (sign of the coefficient β) as the training sets. To select the validated biomarkers whose isoform expression is significantly more predictive than the corresponding overall gene expression we estimated the R^2^ distribution of the isoform- and gene-based models using the bootstrap procedure and compared these distributions using a two-sided Wilcoxon signed rank test.

### Final validation of isoform-based biomarkers (UHN)

To test whether the predictive value of the isoform-based biomarkers validated in GRAY was robust to the use of a different pharmacological assay, we decided to leverage a collection of 84 breast cancer cell lines recently used to investigate gene essentiality in breast cancer molecular subtypes [17]. We selected 14 cell lines in this collection that were readily available and showed extreme expressions of the biomarkers of interest (Supplementary Table 1). Selected cell lines were cultured and screened for their response to three targeted agents: lapatinib, AZD6244 and erlotinib, and one chemotherapy, paclitaxel. We used the sulforhodamine B colorimetric (SRB) proliferation assay [35] in 96--well plates to determine the drug dose--response curves. We subtracted the average phosphate buffer saline (PBS) wells value from all wells and computed the standard deviation and coefficient for each triplicate. Data points with coefficient or standard deviation greater than 0.2 were discarded. All the individual treated well values were normalized to the control well values. We used the *PharmacoGx* [20] package to fit the curves using a logarithmic logistic regression method to estimate the AUC sensitivity values. Raw and processed pharmacological data are available through our *PharmacoGx* platform under the UHNBC PharmacoSet.

### Comparison of isoform expression across patient tumors and healthy tissues

To test whether isoform-based biomarkers are specific to cancerous tissue, we compared their expression distribution across patient tumors and healthy tissues. We downloaded the bam files from The Cancer Genome Atlas (TCGA) [19] and the Genotype-Tissue Expression (GTEx) [36] for patient tumor and healthy tissue RNA-seq profiles, respectively. We reprocessed the data using the Tuxedo protocol [14]. Distribution of isoform expression across sample types is compared using one-sided Wilcoxon rank sum test. The direction of the test was determined by the direction of the biomarker association: for biomarkers associated with drug sensitivity, higher expression in cancer was tested and vice versa.

### Research replicability

The pharmacogenomics data used in this study are publicly available through our *PharmacoGx* platform [20]. Our code and documentation are open-source and publicly available through the RNAseqDrug GitHub repository (<u>github.com/bhklab/RNASeqDrug</u>). A detailed tutorial describing how to run our pipeline and reproduce our analysis results is available the GitHub repository. Our study complies with the guidelines outlined in [37,38].

## RESULTS

We developed a meta-analysis pipeline enabling identification of gene- and isoform-level expression-based biomarkers predictive of sensitivity to 15 drugs (Supplementary Table 1; Supplementary Figure 3) across two large pharmacogenomics studies, namely CCLE and GDSC (Figure 1). CCLE used the CellTiter-Glo (Promega) pharmacological assay, while GDSC used Syto60 (Invitrogen) [21], providing us with the opportunity to discover biomarkers generalizable to multiple measures of drug sensitivities. We identified a large set of statistically significant biomarkers for each drug (14 to 3,480 biomarkers with FDR < 5%; Figure 2A). We observed a significantly larger proportion of isoform-based biomarkers are predictive of drug response (Wilcoxon signed rank test p-value < 10^-5^; Figure 2A). For the majority of genes identified as biomarkers, the highest ranking isoform, but not the overall gene expression, is significantly predictive of drug response (Figure 2B).

### Pre-validation in an independent breast cancer dataset

*In vitro* validation of drug response biomarkers in fully independent datasets has been shown to be challenging [31,39–41]. We therefore sought to assess the predictive value of our most promising isoform biomarkers for eight drugs screened both in our training sets and in the independent breast cancer dataset published by Daemen et al. [3] (referred to as GRAY; Supplementary Figure 5), which used the same pharmacological assay as CCLE. We first selected the significant isoform-based biomarkers in our training set that were predictive in breast cancer cell lines (see Methods). We assessed the predictive value of these biomarker candidates in GRAY and tested whether these isoform biomarkers were significantly more predictive than their corresponding gene expression (Figure 3). The validation success rate ranged from 0% (no validated biomarkers for sorafenib and crizotinib) to 41% validated biomarkers for AZD6244 (Supplementary Table 2). We found that the poor validation rate for crizotinib and sorafenib stems from inconsistency in their pharmacological profiles (Supplementary Figure 6). Based on the number and effect size of biomarker candidates that were significant in GRAY, we selected AZD6244, lapatinib, erlotinib and paclitaxel for further validation.

### Final validation using a different pharmacological assay

To test the robustness of our pre-validated biomarkers we generated a new set of drug sensitivity data combined with the RNA-seq profiles of breast cancer cell lines published by Marcotte et al. [17]. This new pharmacogenomic dataset is referred to as UHN. We screened cell lines with a different pharmacological assay (sulforhodamine B assay; SRB) from those used in the training and pre-validation sets. We first cultured cell lines to check their doubling time in a course of 120 hours (Supplementary Table 3). Only cell lines with a growth rate/doubling time that was amenable to the the 5-day SRB assay as a readout for cytotoxicity were considered for testing in the full 9-dose assay. We then assessed the anti-proliferative effect of cell lines to drugs using SRB assay in 96 well plates in triplicates. All the drug dose-response curves passed our quality controls (see Methods).

Similar to the pre-validation performed in GRAY, we considered an isoformic biomarker to be validated if the linear association between its expression and drug sensitivity is both significant and in the same direction (same coefficient sign in the regression model). This resulted in validation of 3 out of 26, 11 out of 23, 1 out of 4 and 10 out of 31 biomarkers for AZD6244, lapatinib, erlotinib and paclitaxel, respectively (Supplementary Table 2). We selected the most significant isoform for each drug and investigated its exon occupancy and correlation compared with the other isoforms of the same gene (Figure 4; Supplementary Figure 7). The selected TGF-α (ENST00000295400), TNKS1BP1 (ENST00000527207) and DUOX1 (ENST00000389037) isoforms were associated with sensitivity to AZD6244, erlotinib and lapatinib, respectively (Figure 4A-C), while the CPEB4 (ENST00000265085) isoform is associated with lack of sensitivity to paclitaxel (Figure 4D). For TGF-α and DUOX1, the predictive isoform was highly correlated with another isoform of the same gene, sharing similar exon occupancy (Figure 4E,G), while predictive isoform for TNKS1BP1 and CPEB4 present a more specific expression pattern (Figure 4F,H). We compared the expression of the selected isoform biomarkers across patient breast tumors and healthy tissue samples to test whether the biomarkers are tumor-specific (Figure 4I-L), which would facilitate their quantification in future *in vivo* and clinical studies. The TNKS1BP1 isoform was significantly more expressed in tumors compared to healthy tissues (p< 0.001; Figure 4J), while TGFA and DUOX1 isoforms were not (Figure 4I,K). However, for the latter isoform we observed a large tail of tumors yielding higher expression of DUOX1 isoform than any of the healthy breast tissues (Figure 4K), suggesting that these patients may respond to the corresponding therapies. As a biomarker associated with lack of sensitivity, low expression in tumors compared to healthy tissue would favor response, which was actually the case for CEBP4 (p< 0.001; Figure 4L).

## DISCUSSION

Although gene expression represents an important class of biomarkers for prediction of drug response *in vitro* [1–6,18], association between gene isoforms and drug sensitivity has not been well studied despite the critical role of alternative splicing in cancer [7]. Our study is the first to describe a genome-wide meta-analysis of isoform-based biomarker predictive of drug response *in vitro* (Figure 1; Supplementary Table 1). Controlling for the large number of isoforms, we found that significantly more genes had one of their isoforms predictive of response compared to overall gene expression for the vast majority of the drugs (Figure 2A). Importantly only a minority of biomarkers were solely predictive based on their overall gene expression and would have been missed by focusing on isoform expressions (Figure 2B), supporting isoforms as a promising, untapped resource for drug response biomarkers.

Recognizing the challenges involved in biomarker discovery and validation from *in vitro* drug screening data [18,21,30,31,33,39,41–43], we further assessed the predictive value of our newly discovered isoform-based biomarkers for four drugs (AZD6244, lapatinib, erlotinib and paclitaxel) in GRAY, a large independent breast cancer pharmacogenomic dataset (Figure 1 and Supplementary Table 1). As expected given the recognized discrepancies in drug sensitivity profiles between large datasets, we obtained a low validation rate (33-51%; Supplementary Table 2) in our first validation phase, despite the fact that this study used the same pharmacological assay as CCLE to generate their drug sensitivity data (CellTiter Glo; Supplementary Table 1). We found that many of the strongest biomarkers were significantly more predictive of drug sensitivity at the isoform level compared to the overall gene expression level (Wilcoxon signed rank test p< 0.05; Figure 3).

Given that we and others have shown that the choice of pharmacological assay strongly influences drug sensitivity measurements [18,21,30], we sought to validate our candidate isoform biomarkers using the sulforhodamine B assay (SRB), which differs from the assays used in the training and pre-validation datasets (Figure 1). We selected 14 breast cancer cell lines and screened them with the set of four drugs. Despite the small sample size, we validated 10 isoform biomarkers (p< 0.05; Supplementary Table 2). We selected the most predictive isoform for each drug to investigate its correlation with the other isoforms of the same gene and its distribution across patient tumor and healthy tissue samples (Figure 4). As a biomarker predictive of response to the MEK inhibitor AZD6244 in breast cancer, we identified ENST00000295400, one of the longest isoforms of the transforming growth factor alpha (TGF-α), which codes a protein with 160 amino acids. The expression of this isoform is highly correlated with ENST0000041833 which has a very similar transcriptomic structure (Figure 4E) and codes for a protein with just 4 less amino acids. However the other seven isoforms of TGF-α are poorly correlated with ENST00000295400 (ρ<0.8) and the inclusion of the extra exons resulted in the loss of predictive value for TGF-α overall expression. TGF-α is a member of the epidermal growth factor (EGF) family, which binds to the EGF receptors (EGFR) on cell surface and activate a signalling pathway for multiple cell proliferation events including the MAPK/ERK pathway involved in cell proliferation [44,45]. It has been shown that increased TGF-α expression causes persistent stimulation of the EGFR by creating an autocrine feedback loop [45]. The association between ENST00000295400 expression and response to MEK inhibition suggests that this feedback loop may make the breast cancer cells reliant on activated MAPK/ERK pathway and consequently increase their sensitivity to AZD6244.

We investigated the association between isoform expressions and sensitivity to lapatinib, a dual tyrosine kinase inhibitor which interrupts the HER2/neu and epidermal growth factor receptor (EGFR) pathways. Concurring with the literature [46], we found that breast cancer cell lines overexpressing ERBB2 were highly sensitive to lapatinib (Figure 3B). However, this biomarker is not isoform-specific as overall ERBB2 expression is similarly predictive of drug response (Supplementary Figure 8). We further identified ENST00000527207, the shortest protein-coding isoform for TNKS1BP1 as the strongest isoform-specific biomarker (Figure 4B). No other TNKS1BP1 isoforms are strongly correlated with ENST00000527207 (ρ< 0.8), supporting its unique predictive value compared to overall expression (Figure 4F). TNKS1BP1 was originally identified as an interaction protein of tankyrase 1, which belongs to the poly(ADP-ribose) polymerase (PARP) superfamily; however its function is poorly characterized. Although TNKS1BP1 association with drug response is intriguing, the dominent predictor of response will remain ERBB2 expression in clinical setting.

Our results indicate that sensitivity to the EGFR inhibitor, erlotinib, can be predicted by the expression of the ENST00000389037 isoform of DUOX1 (Figure 4C). This isoform was highly correlated with ENST00000321429, which differs only by a single splicing event, but was not strongly correlated with the other 10 isoforms (ρ < 0.8; Figure 4G). DUOX1 has been shown to induce ATP-mediated EGFR transactivation in airway epithelial cells [49] and more recently in squamous-cell cancer [50]. Although there is no evidence yet for EGFR transactivation in breast cancer, the association between DUOX1 and erlotinib sensitivity suggests that breast cancer cell lines overexpressing DUOX1 may be reliant on activated EGFR signaling for survival, making them more vulnerable to EGFR inhibition. Given evidence for some clinical activity of EGFR inhibitors in breast cancer, our result uncovers new opportunities to characterize this pathway towards the development of biomarker driven treatment strategies for this class of drugs.

Lack of sensitivity or innate resistance to chemotherapies is a major issue in current breast cancer management [51]. Our results indicate that the expression of the ENST00000265085 isoform of the cytoplasmic polyadenylation element binding protein 4 (CPEB4) genes is associated with lack of sensitivity to paclitaxel in breast cancer cell lines (Figure 4D). None of the remaining nine CPEB4 isoforms is highly correlated with ENST00000265085 (ρ<0.8; Figure 4H). The cytoplasmic polyadenylation element binding proteins combine a sequence-specific RNA-binding protein with a RNA-recognition motif and a zinc-finger [52,53] and associate with specific sequences in mRNA 3ʹ untranslated regions to promote translation [54]. Elevated CPEB4 expression have been associated with tumor growth, vascularization, migration, invasion, and metastasis in multiple cancer types [55–58]. Xu and Liu found that the CPEB4 targeted genes, such as BIRC5 [59] and IGF2 [60], are related to chemotherapy resistance and suggested CPEB4 as a marker of resistance to paclitaxel and cisplatin [56]. These mechanistic studies are consistent with our finding that the expression of the first isoform of CEBP4 correlates with lack of sensitivity to paclitaxel; additional characterization of the biology underlying the isoform specificity of this association would be of substantial interest (Figure 4D).

This study has several potential limitations. First, our biomarker discovery pipeline is restricted to univariate linear association between gene and isoform expression and drug sensitivity. These two restrictions have been imposed to mitigate the risk of overfitting as the development of multivariate, potentially nonlinear predictors of *in vitro* drug sensitivity has been proven to be challenging [31,39]. Larger sample size of compendia of pharmacogenomic datasets will be necessary to overcome this. A second limitation lies in the use of a single processing pipeline to quantify expression of each individual transcripts from Illumina RNA-seq data. We choose to use the Tuxedo protocol for RNA-seq [14] because it is one of the most widely-used suite of tools for transcript expression analysis. We recognize that many alternatives exist [61–63] but their comparison is out of the scope of the present study. Third, the validation of our biomarkers is limited to breast cancer cell lines, the only tissue type for which we had independent pharmacological and molecular data. The release of additional large-scale pharmacogenomic datasets will enable validation in more tissue types, to which our computational approach can readily be applied. Lastly, we are aware that our comparison of the tumour and healthy tissue expression profiles extracted from the TCGA and GTEx projects, respectively, might be biased due to the inevitable batch effects and other technical variations across laboratories. To alleviate this issue, the TCGA and GTEx RNA-seq raw data have been downloaded and reprocessed using the same analysis pipeline to ensure that the transcript expression values are comparable.

## CONCLUSION

The advent of RNA-sequencing technology enables efficient quantification of alternatively-spliced transcripts in cancer cells. Our genome-wide search for biomarkers demonstrates that gene isoforms consitute a rich resouce of transcriptomic features associated with response to targeted and chemotherapies *in vitro*. Our results suggest that isoform-based biomarkers are more frequent and more significantly associated with drug sensitivity than overall gene expression, opening new avenues for future biomarker discovery for *in vitro* and *in vivo* drug screening.

## ACKNOWLEDGEMENTS

The authors would like to thank the investigators of the Genomics of Drug Sensitivity in Cancer (GDSC), the Cancer Cell Line Encyclopedia (CCLE), Drs. Joe W. Gray and Benjamin G. Neel who have made their invaluable data available to the scientific community.

## FUNDING

Z Safikhani was supported by the Cancer Research Society (Canada; grant #19271) and the Ontario Institute for Cancer Research through funding provided by the Government of Ontario. P Smirnov was supported by the Canadian Cancer Society Research Institute. B Haibe-Kains was supported by the Gattuso Slaight Personalized Cancer Medicine Fund at Princess Margaret Cancer Centre, the Canadian Institute of Health Research (grant #340176) and Natural Sciences and Engineering Research Council (grant #357163).

